# Efficacy of land use designation in protecting habitat in the miombo woodlands: Insights from Tanzania

**DOI:** 10.1101/117622

**Authors:** Alex L. Lobora, Cuthbert L. Nahonyo, Linus K. Munishi, Tim Caro, Charles Foley, Colin M. Beale

## Abstract

Loss of natural landscapes surrounding major conservation areas compromise their future and threaten long-term conservation. We evaluate the effectiveness of fully and lesser protected areas within Katavi-Rukwa and Ruaha-Rungwa ecosystems in south-western Tanzania to protecting natural landscapes within their boundaries over the past four decades. Using a time series of Landsat satellite imageries of September 1972, July 1990 and September 2015, we assess the extent to which natural habitat has been lost within and around these areas mainly through anthropogenic activities. We also test the viability of the remaining natural habitat to provide connectivity between the two ecosystems. Our analysis reveals that while fully protected areas remained intact over the past four decades, lesser protected areas lost a combined total area of about 5,984 km^2^ during that period which is about 17.5% of habitat available in 1972. We also find that about 3,380 km^2^ of natural habitat is still available for connectivity between the two ecosystems through Piti East and Rungwa South Open Areas. We recommend relevant authorities to establish conservation friendly village land use plans in all villages surrounding and between the two ecosystems to ensure long-term conservation of these ecosystems.

## 1. Introduction

About 15% of the land worldwide is currently designated as protected areas (henceforth “PAs”) for biodiversity conservation (Juffe-Bignoli *et al.,* 2014), and efforts are underway to further extend this to 17% of all terrestrial land and 10% of coastal and marine areas by 2020 (Di Minin & Toivonen, 2015). These efforts indicate the importance that countries and the international communities attach to the role PAs play in preserving natural landscapes and reducing biodiversity loss over long-term (Dudley, 2008; Serra *et al.,* 2008; Wyman & Stein, 2010; Schulz *et al.,* 2010; Gao and Liu, 2010; Gibbs *et al.,* 2010; Lung & Schaab, 2010; Leroux *et al.,* 2010; Azadi & Hasfiati, 2011; Estes *et al.,* 2012; Bailey *et al.,* 2016). However, poverty, population pressures and escalating demand for natural resources compounded by conflicting national policies, poor governance and weak institutions (Rao *et al.,* 2009), have influenced the ability of PAs to fulfil their role (Brashares *et al.,* 2001; Dwivedi *et al.,* 2005; Shalaby & Tateishi, 2007; Bakr *et al.,* 2010). While the expansion of PAs coverage and their contribution to nature conservation is well recognized (Pimm *et al.,* 1995; Baillie *et al.,* 1996; Myers *et al.,* 2000; Bruner *et al,* 2001; Rodrigues *et al.,* 2004a,b; Jenkins & Joppa, 2009; Leverington *et al.,* 2010; Machumu & Yakupitiyage, 2013), there is increasing argument surrounding PAs’ effectiveness in protecting biodiversity loss due to the wide spreading anthropogenic activities within and outside their boundaries (Pimm & Raven, 2000; DeFries *et al.,* 2005; Stoner *et al.,* 2007a; Gardner *et al,* 2007; Bradshaw *et al.,* 2009; Ahrends *et al.,* 2010; Gibson *et al.,* 2011; Laurance *et al.,* 2011, 2012; Ghimire & Pimbert, 2013; Clark *et al.,* 2013; WWF, 2016).

Tanzania maintains a variety of PA categories which allow different levels of legal restrictions on resource use including fully protected areas (henceforth “FPAs”) (constituting 17% of land surface) comprising of national parks (only allow photographic tourism), game reserves (permit tourist hunting), forest reserves (some permit selective logging) and the Ngorongoro Conservation Area (NCA) which is similar to national parks but allows cattle grazing by indigenous Maasai pastoralists (MNRT, 2007; Stoner *et al.,* 2007a). The rest of the PAs (constituting 18% of the country’s land surface) are considered as lesser protected (henceforth “LPAs”) and include Game Controlled Areas (GCAs) and Open Areas (OAs) where extractive resource use is permitted under license (MNRT, 2007). Wildlife Management Areas (WMAs) comprise Tanzania’s newest protection category that aims to promote wildlife management at the village level by allowing rural communities and private land holders to manage wildlife on their land for their own benefit and devolving management responsibility of the settled areas and areas outside unsettled PAs to rural people and the private sector (USAID, 2013; WWF, 2014).

Despite having a variety of PA categories in place, there has been little effective evaluation of how well the different classes actually prevent habitat loss although efforts have been made on assessing wildlife numbers in these areas (Gardner *et al.,* 2007; Stoner *et al.,* 2007b). There is some evidence that overall habitat degradation is lower in Tanzania’s PAs than outside (Pelkey *et al.,* 2000; Beale *et al* 2013), but effective protection of forests within protected areas is certainly mixed (Pfeifer *et al.,* 2012).While the IUCN guidelines advocate a rule of thumb law enforcement effort of one ranger/scout for every 10-50 sq.km (David *et al.,* 2016), the mean effort available for FPAs and LPAs within the study area is one scout/ranger for every 143 and 346 sq.km respectively indicating minimal enforcement in both categories (Nahonyo, 2005). Here, we use time series satellite datasets to investigate the status of land-cover in the various landuse designations in the study area over the past four decades. More specifically, we assess the spatial extent of deforestation in both FPAs and LPAs during that period, establish which of the two PA categories is effective in halting habitat change during that period and quantify available potential habitat for connectivity between the two ecosystems.

## 2.0 Study area, materials and methods

### 2.1 Study area

The study area covers about 109,050 km^2^ and lies between latitude 6^0^15'59.38" to 8^0^10'23.78" S and longitude 30^0^45'13.29" to 35^0^28'34.44" E. It comprises Katavi-Rukwa ecosystem in the west, a contingent of Game Reserves (henceforth “GRs”), GCAs and OAs in the central part as well as Ruaha-Rungwa ecosystem in the East (Figure 1). About 45,961 km^2^ of this area is designated as FPAs (2 NPs, 7 GRs), 34,196 km^2^ designated as LPAs (8 GCAs and 8 OAs) herein also referred to in our analysis as Region Of Interest (ROI). A further 28,893 km^2^ of land within the study area was considered as unclassified land and hence excluded from the analysis and includes towns and highly populated hamlets north and south of Katavi National Park, east of Muhesi Game Reserve and south of Lunda - Nkwambi GCA (Figure 1).

**Figure 1:**
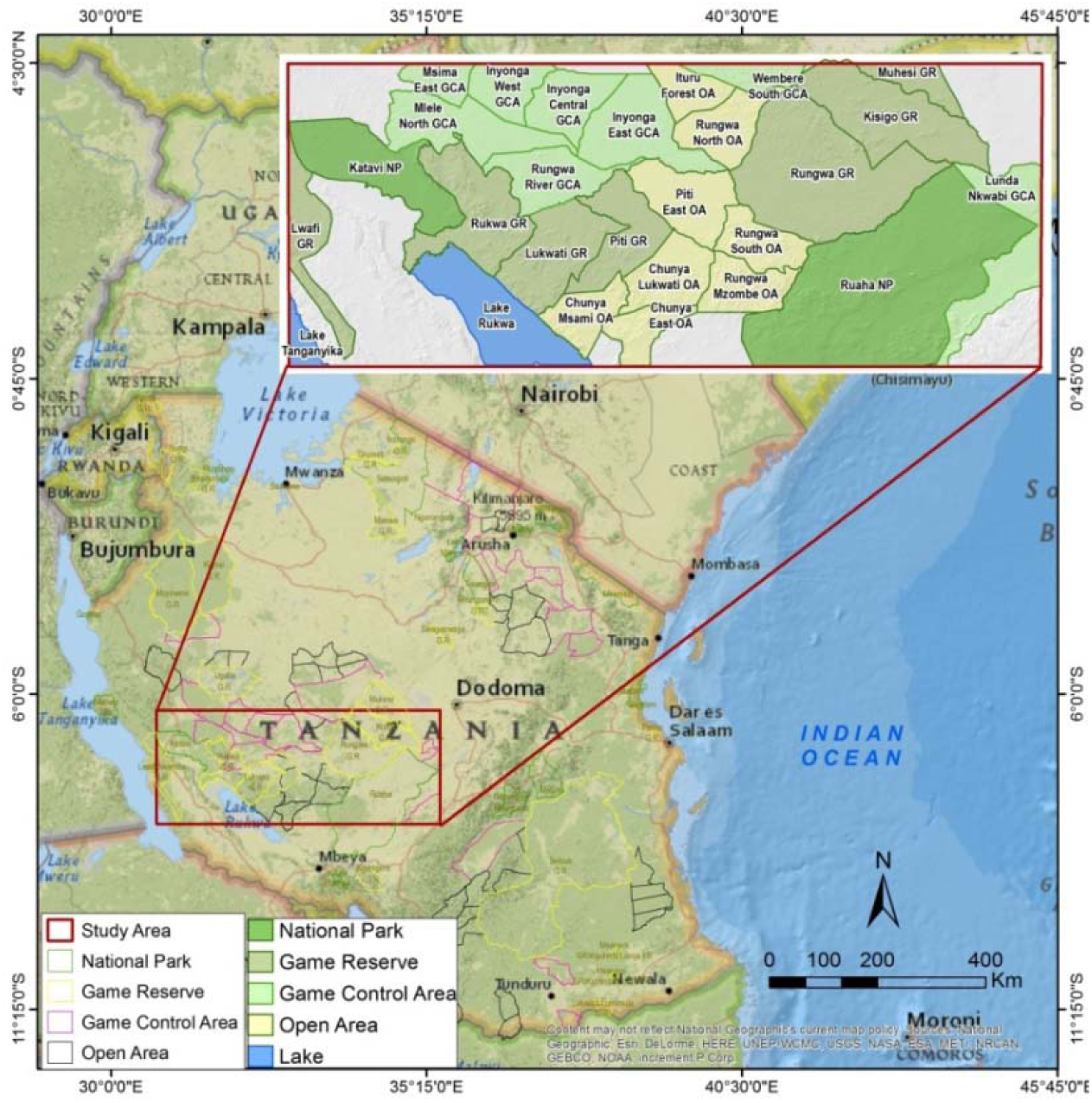
Study Area.

### 2.2 Method

#### 2.2.1 Data collection

Our analysis employed Landsat Multispectral Scanner (MSS), Landsat Thematic Mapper (TM) and Landsat Enhanced Thematic Mapper (ETM^+^) for 1972, 1990 and 2015 respectively. The study area contained an intersection of 12 Landsat footprints (path/row 168/064, 169/064, 170/064, 171/064, 171/065, 170/065, 169/065, 168/065, 171/066, 170/066, 169/066 (Annexes 1-3). Cloud free Landsat images were downloaded from USGS website in single band GEOTIFF format pre-processed for atmospheric correction, geometric correction and noise-removal. The TM sensor has seven spectral bands (Boettinger *et al.,* 2008; Brink & Eva, 2009) and is primarily designed to detect reflected radiation from the Earth’s surface in the visible and near-infrared (IR) wavelengths (Shalaby & Tateishi 2007).

We obtained land-use data from the Tanzania Wildlife Research Institute (TAWIRI), a local government organ responsible for providing scientific information for promoting the development, improvement and protection of the wildlife industry. The dataset included boundaries of NPs, GRs, GCAs, and OAs and road networks.

#### 2.2.2 Image pre-processing

We used a combination of ENVI version 5.1 (Exelis Visual Information Solutions, Boulder, Colorado, USA) and ArcMap module of the ArcGIS 10x software for image preparation and processing. The first step involved geo-referencing images using known locations taken across the study area to reduce registration error (Jensen, 1996). We then applied image enhancement tool in ArcMap to improve visual interpretability by increasing the apparent distinction between different image features (Bradley & Mustard, 2005; Shalaby & Tateishi 2007). We further normalised each band stretching from 0 to 255 to improve visibility of different bodies with similar tones. To improve interpretability, we colour composited individual image bands to generate Colour Composites. For analysis we used bands 2, 3, and 4 for Landsat TM and ETM+ images and bands 6, 5 and 4 for Landsat MSS images.

#### 2.2.3 Image processing

Image interpretation involved a combination of supervised and unsupervised (hybrid) classification (Brink & Eva, 2009). We first performed unsupervised classification using an Iterative Self-Organizing Data Analysis Technique Algorithm (ISODATA) which has been shown to perform better because of the statistical power it employs when classifying an unknown pixel (Shalaby & Tateishi, 2007; Brink and Eva, 2009). We set our preliminary classification result to yield a maximum of 30 spectral classes for historical (1972, 1990) and current (2015) maps (Ball & Hall, 1965). To obtain hybrid land cover maps, we visually interpreted and assigned the relevant landcover classes (from unsupervised classification) with the help of field data, Google earth images, expert knowledge and the recent countrywide landcover map developed by the National Forest Resources Monitoring and Assessment (NAFORMA) project (NAFORMA, 2015). Landcover classes for the final maps included Closed woodland, Open woodland, Bushland, Settlement/Cultivation, Wetland and Water. Description of theses land cover classes are provided in Table 1.

**Table 1:** Description of different land covers classes of the study area

| Land cover | Description |
| --- | --- |
| Closed woodland | Tree layer with crown cover >40% |
| Open woodland | Tree layer with crown cover $\geq 10\%$ and $\leq 40\%$ |
| Bushland | Mixed vegetation types including thicket, dense bushland, bushland with scattered cultivation and Open bushland |
| Settlement/Cultivation | Open and cultivated agricultural grasslands mixed with settlement, grassland, shrubland, and impervious surfaces |
| Wetland | Vegetated lands with a high water table and inundated vegetation |
| Bare/Open Areas | Land areas of exposed soil surface as influenced by human impacts and/or natural causes. It contains sparse vegetation with very low plant cover value as a result of overgrazing, woodcutting, etc. |
| Water | Areas covered with water most of the year, Lakes, ocean, river, etc. |

Source: National Forestry Resources Monitoring and Assessment of Tanzania Field Manual (NAFORMA, 2010).

#### 2.2.4 Accuracy assessment

We assessed accuracy (Landgrebe, 2003; Mather, 2004) using 254 random points chosen to represent different land cover classes across the study area collected in 2015. We determined both omission error/producer accuracy to measure how well our images have been classified, commission error/user’s accuracy to determine reliability of a pixel class on the map and the category on the ground as well as Kappa (Hat) to measures of agreement between the classification and the reference data (Olofsson *et al.,* 2014). Accuracy above 85% is considered within the acceptable range (Anderson *et al.,* 1976; Lins & Kleckner, 1996) while Kappa statistic above 80% is considered strong (Jensen, 1996). To obtain overall accuracy, we divided the total number of correct pixels (diagonal) by the total number of pixels in the error matrix (Olofsson *et al.,* 2014). We could not assess accuracies for historical land-cover maps due to the lack of historical reference points.

#### 2.2.5 Detecting change

Change detection (Singh 1989; Coppin, 2004) involved identifying pixels that were previously natural habitat in 1972 but later changed to field crops and settlements in 2015. To do this we carried out a pixel based raster analysis in R (R Core team, 2016) in three phases namely between 1972 and 1990, 1990 and 2015 as well as the overall change between 1972 and 2015 (Appendix H). The analysis generated three different change tables as per the above phases and subsequently merged and used to calculate respective changes.

## 3. Results

### 3.1 Land-cover classification and accuracy assessment

Overall accuracy assessment and Kappa coefficients for the 2015 final land-cover map (Figure 2) are 89% and 87% respectively (Table 2). User’s and producer’s accuracies for individual land-cover classes are high suggesting correct assignment of individual classes during classification (Table 2).

**Figure 2:**
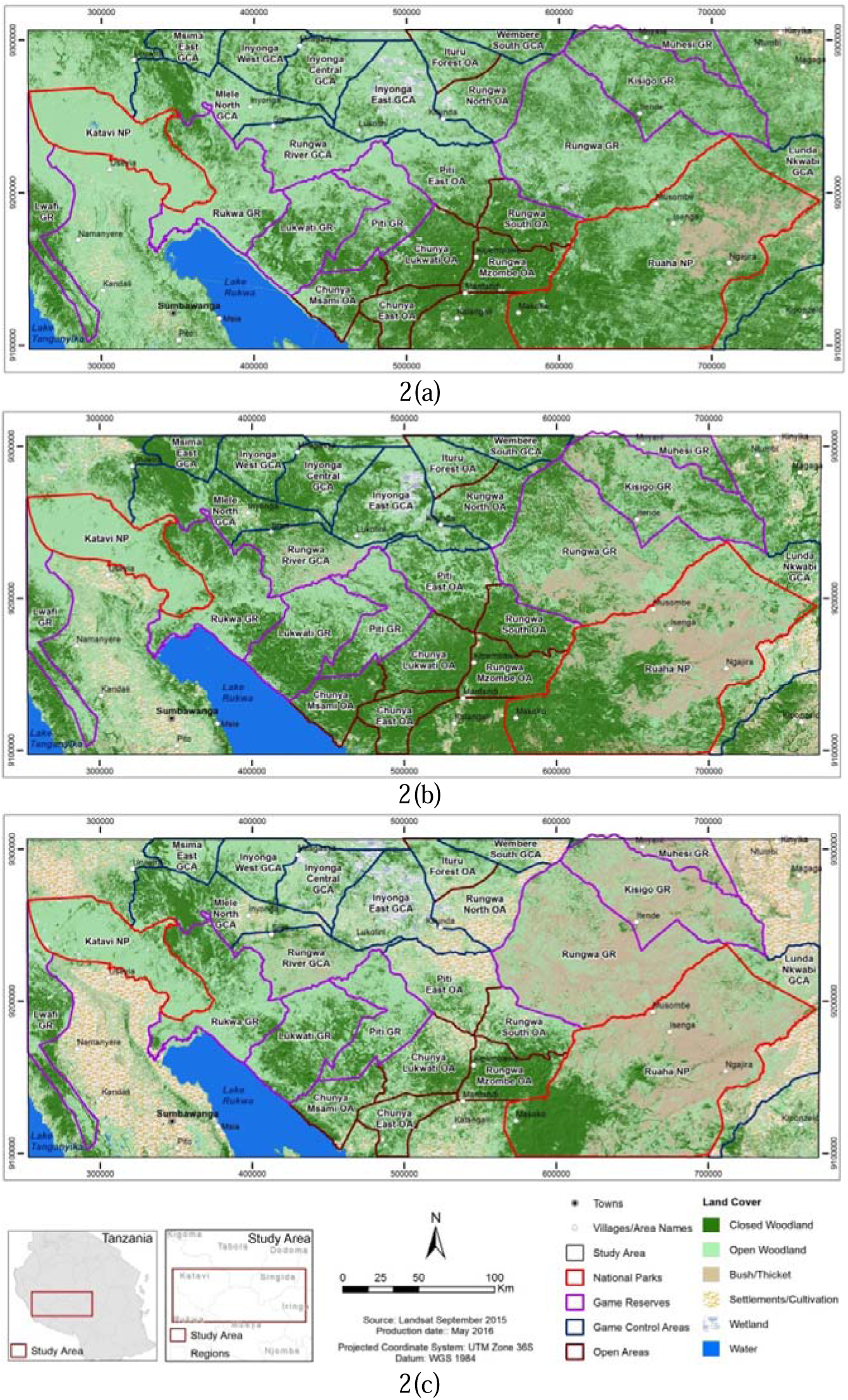
Landcover maps for the study area showing trends in habitat decline over years. 2 (a) Landcover map of the study area in 1972 indicating that closed and open woodlands combined comprised a large proportion of the study area then. 2(b) Landcover map for the study area in 1990 indicating dominance of closed and open woodlands combined still comprised a large proportion of the study area. 2(c) Landcover map of the study area in 2015 indicating massive reduction in natural habitat. National Parks (Katavi and Ruaha) and Game Reserves (Rukwa, Lukwati, Piti, Rungwa, Kisigo and Muhesi) landscapes remained intact whilst Game Controlled Areas (GCAs and OAs) are significantly reduced to crop fields and settlements at the expense of natural habitat.

**Table 2:** Classification error matrix based on ground truth data collected in the study area

| Classification data | Reference data |  |  |  |  |  | User accuracy |
| --- | --- | --- | --- | --- | --- | --- | --- |
|  | Closed Woodland | Open Woodland | Bushland | Crop fields | Wetland | Water |  |
| Closed woodland | 39 | 1 | 0 | 0 | 0 | 0 | **98** |
| Open woodland | 11 | 49 | 6 | 2 | 2 | 0 | **70** |
| Bushland | 0 | 0 | 43 | 2 | 2 | 0 | **91** |
| Crop fields | 0 | 0 | 0 | 46 | 0 | 0 | **100** |
| Wetland | 0 | 0 | 1 | 0 | 15 | 0 | **94** |
| Water | 0 | 0 | 0 | 0 | 0 | 35 | **100** |
| **Producer Accuracy** | **78** | **98** | **86** | **92** | **79** | **100** |  |
| Overall Accuracy | 89 |  |  |  |  |  |  |
| Kappa statistics | 87 |  |  |  |  |  |  |

Note: Percentage of pixels classified is shown by class with kappa statistic and producer, user and overall accuracy.

### 3.2 Status of landuse/cover in and around LPAs (GCAs and OAs)

In 1972, areas between and surrounding the two ecosystems entirely comprised of natural habitat (Figure 2a). Two decades later (between 1972 & 1990), these areas combined lost an estimated 1,060 km^2^ of natural habitat to crop fields and settlements equivalent to 3% of natural habitat available in 1972 (Figure 2b, Appendix F). Much of the loss occurred between 1990 and 2015 where an estimated 4,900 km^2^ of natural habitat was lost to crop fields and settlements which is equivalent to 6% of habitat available in 1972 (Figure 3). Piti East and Rungwa South Open Areas which provide the potential habitat for connectivity between the two ecosystems lost a combined total area of about 1,200 km^2^ of natural habitat to crop field and settlement between 1972 and 2015, with 89% of the loss occurring between 1990 and 2015 (Figure 2c). Overall, habitat within and around LPAs have lost an estimated 5,984 km^2^ (equivalent to 17.5%) of natural habitat to agricultural and settlement activities between 1972 and 2015 (Figure 3, Appendix F).

**Figure 3:**
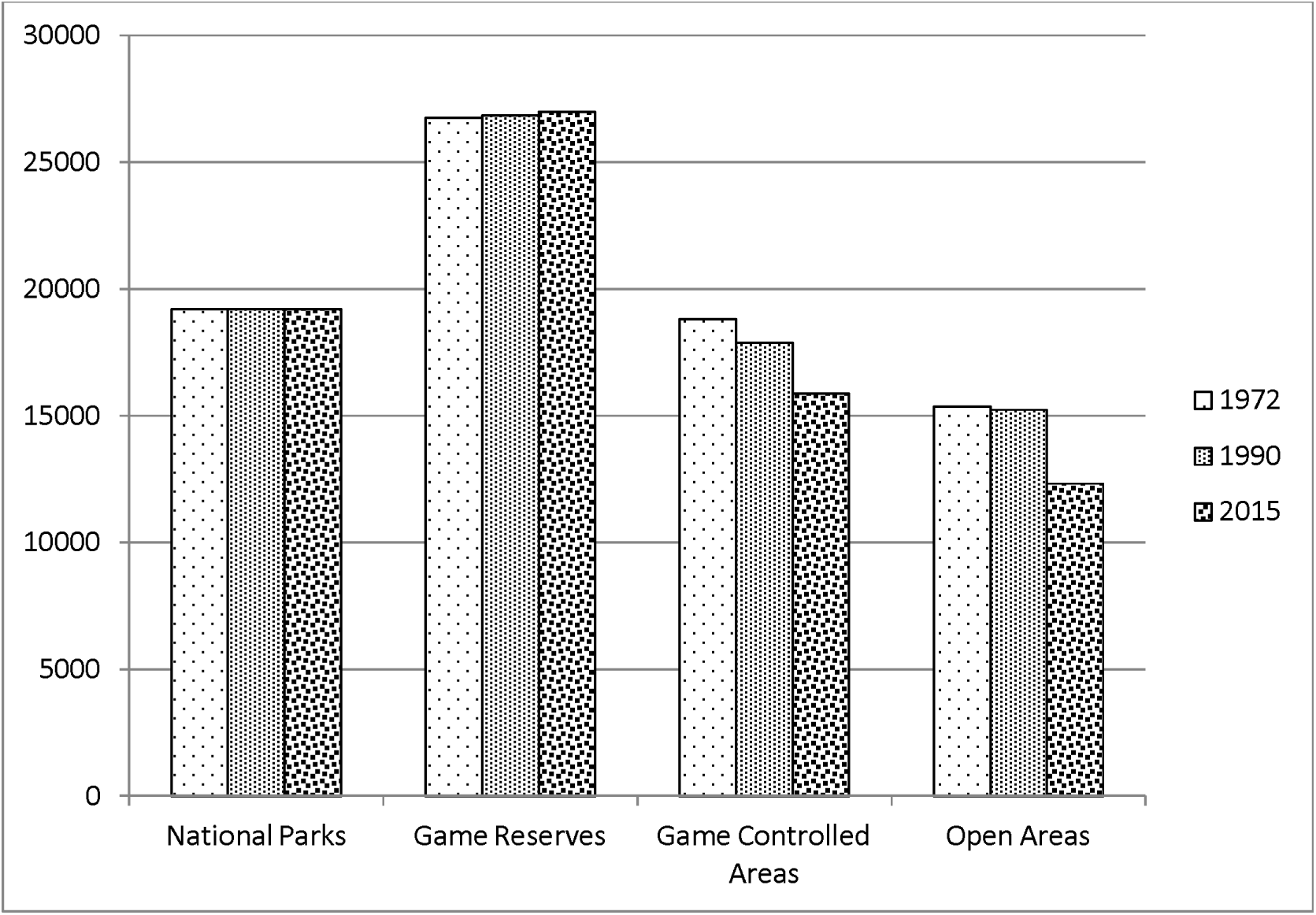
Overall habitat reduction (in km^2^) in all four land use categories in the study area between 1972 and 2015. Habitat in National Parks and Game Reserves remain unchanged throughout suggesting the effectiveness of full protection.

### 3.3 Status of landuse/cover in FPAs (NPs and GRs)

Apart from subtle habitat losses detected in Muhesi and Rukwa Game Reserves, FPAs remained relatively intact throughout suggesting their land use designation has been effective in preventing habitat loss over long term (Figure 3, Appendix E).

## 4. Discussion

Our analysis revealed that FPAs are effective at preventing total habitat loss in the study area, despite minimal investment in enforcement (Figure 3, Appendix E). Our results support previous findings conducted across the country which found an increase in vegetative cover in FPAs with LPAs suffering worse habitat degradation than areas with no legal protection (Pelkey *et al.,* 2000). In addition, a study conducted across tropical countries that found 83% of PAs were effective at preventing land clearance (Bruner *et al.,* 2001). While habitat within FPAs in the study area remained relatively intact with minimum investment, we note that this may overlook losses of high valuable tree species such as African Blackwood *Dalbergia melanoxylon* and sealing-wax tree *Pterocarpus angolensis* (Caro *et al.,* 2005) that may be selectively removed from within the miombo woodlands without total habitat loss (Appendix G). Our on the ground experience affirms this is happening in this part of Tanzania on large scale (Caro *et al.,* 2005). Extensive selective cutting of high valued tree species is not only widespread in all PA categories within the study area but also documented elsewhere within FPAs in central Tanzania where increasing rarity has compelled wood carvers to shift to less preferred Miombo species *Brachystegia speciformis* (Madulu, 2001). Similarly, there is evidence of widespread elephant poaching within FPAs across the sub Saharan Africa (Brennan & Kalsi, 2015), and in the Ruaha ecosystem in particular (TAWIRI, 2014;Chase *et al.,* 2016), suggesting again a distinction between successful habitat preservation and successful protection of high value species within that habitat.

In contrast to FPAs, LPAs in the ROI have experienced rapid habitat loss in the study area, with 17.5% transformed to agricultural activities between 1972 and 2015. Most LPAs within the study area and elsewhere in the country have limited resources for enforcement. LPAs such as Piti East and Rungwa South OAs rely for help from adjacent FPAs namely Piti and Rungwa Game Reserves respectively (pers. obs.). Much of the conversion in these areas can be attributed to weak enforcement coupled with increased human population densities attracted by nearby resources (Wittemyer *et al.,* 2008; c.f. Serengeti NP: Metzger *et al.,* 2010) and the poor ability of the land to sustain agriculture (Campbell, 1996; Malmer, 2007). In many cases, human conversion of natural habitats occur most rapidly in the locations important to biodiversity because humans and biodiversity tend to be concentrated in the same locations (Hansen *et al.,* 2012), often resulting in elevated levels of exploitation in these locations (Brashares *et al.,* 2001; Parks & Harcourt, 2002).

Miombo woodland generally occurs on low fertility soils which limits intensive agriculture, causing people to adopt destructive and unstable forms of agriculture and pastoralism activities (MacKinnon *et al.,* 1986; Malmer, 2007), often at the expense of natural habitat (Mwalyosi, 1991). This leads to shifting agriculture not only because fertility declines fast on newly-cleared fields, but also because invasion of weeds makes it labour intensive to re-cultivate and hence they are often abandoned (MacKinnon *et al.,* 1986). Such losses outside FPAs are a concern, because wide ranging species such as African elephants require vast areas that may not be provided by FPAs to guarantee long-term survival (McNeely & Scherr, 2003; Zuidema & Sayer 2003; Laurance *et al.* 2006; Michalski *et al.* 2007; Vandermeer & Perfecto 2007; Harvey *et al.* 2008; Hansen *et* al.,2012). For example, African elephants spend a substantial amount of their time outside protected areas in Northern Tanzania (e.g. Tarangire National Park:Galanti *et al.,* 2006) and elsewhere (Douglas-Hamilton *et al.,* 2005). Escalating habitat loss through expanding agricultural and pastoralist activities within LPAs and around FPAs across the country threatens the few remaining corridors (Caro *et al.,* 2009; Jones *et al.,* 2009) and could potentially reduce genetic exchange (Rodrigues *et al.,* 2004a; Newmark, 2008; Graham *et al.,* 2009;Craigie *et al.,* 2010).

Currently, Tanzania has about 17% of her land area devoted to wildlife conservation in PAs where no human settlement is allowed and further 18% of its surface area to PAs where wildlife co-exist with humans (MNRT, 2007). This network of PAs is coming under intense public scrutiny, with the government seeking to ensure development and conservation are adequately balanced across the country (Rutasitara *et al.,* 2010) and there is little desire to increase the proportion of land in FPAs (Kideghesho *et al.,* 2007). Indeed, the recent Southern Agricultural Growth Corridor of Tanzania (SAGCOT) initiative 2011 is explicitly targeting intensification of agriculture within the coastal plains and the valleys of Kilombero and Ruaha, on the hills and valleys of the Southern Highlands and the Usangu flats, including many areas of high biodiversity value (Milder *et al,* 2013; Wuyts & Kilama, 2015).

Out of an estimated 26,090 km^2^ of natural habitat that remains in ROI today, about 3,380 km^2^ within Piti East and Rungwa provides potential habitat for connectivity between Rukwa-Katavi and Ruaha-Rungwa ecosystems (Appendix D). Both Piti East and Rungwa permit utilization through hunting of big game animals. However, Piti East OA was recently abandoned by hunting companies due to increasing number of cattle in the vicinity coupled with a substantial decline in large mammal populations, rendering the block currently uneconomical. The lack of interested hunting companies to invest in Piti East OA makes it vulnerable to greater habitat loss. Adequately protecting these two areas could secure connectivity between the two largest populations of African elephants of Katavi-Rukwa and Ruaha-Rungwa ecosystems in south-western Tanzania (Jones *et al.,* 2009).

## 3. Recommendations

Protecting the remaining habitat connecting Ruaha-Rungwa & Katavi-Rukwa ecosystems is a high priority. Increasing enforcement in LPAs within the study area and the country at large could help reduce the on-going conversions given the current political situation encouraging WMA establishment than new FPAs, although in practice it achieves the same result if well implemented. Additionally, the land here is marginal at best for agriculture, so the development of a WMA offers solution that could allow greater local community buy-in. The area will need restoration before it can be economically self-sustaining and this needs investment, with some initial funds expected from Wildlife Conservation Society (WCS), Reducing Emissions from Deforestation and Degradation (REDD) programs offers others funding sources. Our analysis suggests that without increased protection this corridor will be lost in the next few years.

## Acknowledgments

We would like to thank Ms Rose Mayienda for assisting with data processing, Wildlife Conservation Society (WCS) Tanzania country office and the Tanzania Wildlife Research Institute (TAWIRI) for financially supporting this study.

## Funding

This research did not receive any specific grant from funding agencies in the public, commercial, or not-for-profit sectors.

## Appendices

## Appendix A: List of satellite images used in the study

| Sensor type | Scene No. | Date of acquisition |
| --- | --- | --- |
| MSS | LM11800641974007AAA04 | 07/01/1974 |
| MSS | LM11800651974007AAA04 | 07/01/1974 |
| MSS | LM11800661972216AAA02 | 03/08/1972 |
| MSS | LM11810641973247AAA02 | 09/09/1972 |
| MSS | LM11810651972253AAA05 | 09/09/1972 |
| MSS | LM11810661973265AAA05 | 09/09/1972 |
| MSS | LM11820641973176FAK03 | 09/10/1972 |
| MSS | LM11820651972272AAA05 | 09/10/1972 |
| MSS | LM11820661972254AAA05 | 09/10/1972 |
| MSS | LM11830641973231AAA05 | 19/08/1973 |
| MSS | LM11830651972255AAA05 | 11/09/1972 |
| MSS | LM11830661972255AAA05 | 11/09/1972 |
| TM | LE71680641999305EDC00 | 07/11/1990 |
| TM | LT51680651999041JSA00 | 07/11/1990 |
| TM | LT51680661998134JSA00 | 07/11/1990 |
| TM | LT51690641997218JSA00 | 06/16/1990 |
| TM | LT51690651997186JSA00 | 06/16/1990 |
| TM | LT51690661998285JSA00 | 06/16/1990 |
| TM | LT51700641997177JSA00 | 23/06/1990 |
| TM | LT51700651997177JSA00 | 23/06/1990 |
| TM | LT51700661997177JSA00 | 23/06/1990 |
| TM | LT51710641997184JSA00 | 16/07/1990 |
| TM | LT51710651997232JSA00 | 16/07/1990 |
| TM | LT51710661997232JSA00 | 16/07/1990 |
| ETM+ | LC81680642014034LGN00 | 09/18/2015 |
| ETM+ | LC81680652014034LGN00 | 09/18/2015 |
| ETM+ | LC81680662013303LGN00 | 09/21/2015 |
| ETM+ | LC81690642013230LGN00 | 09/09/2015 |
| ETM+ | LC81690652013230LGN00 | 09/09/2015 |
| ETM+ | LC81690662013294LGN00 | 09/09/2015 |
| ETM+ | LC81700642013253LGN00 | 08/31/2015 |
| ETM+ | LC81700652013253LGN00 | 08/31/2015 |
| ETM+ | LC81700662013269LGN00 | 08/31/2015 |
| ETM+ | LC81710642013260LGN00 | 09/23/2015 |
| ETM+ | LC81710652013244LGN00 | 09/23/2015 |
| ETM+ | LC81710662013228LGN00 | 09/23/2015 |

## Appendix B: Landsat MSS satellite index

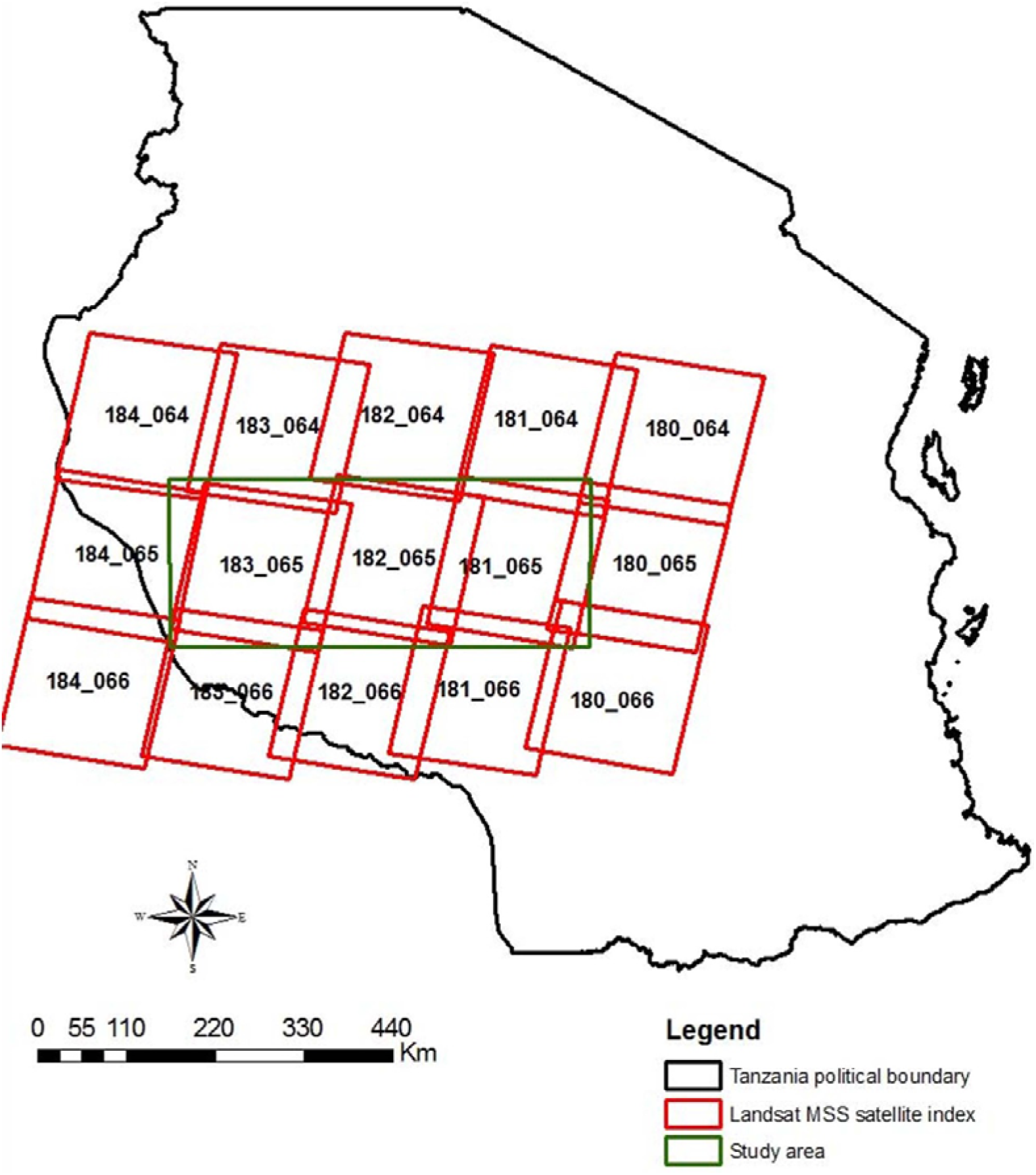

## Appendix C: Landsat TM/ETM+ satellite index

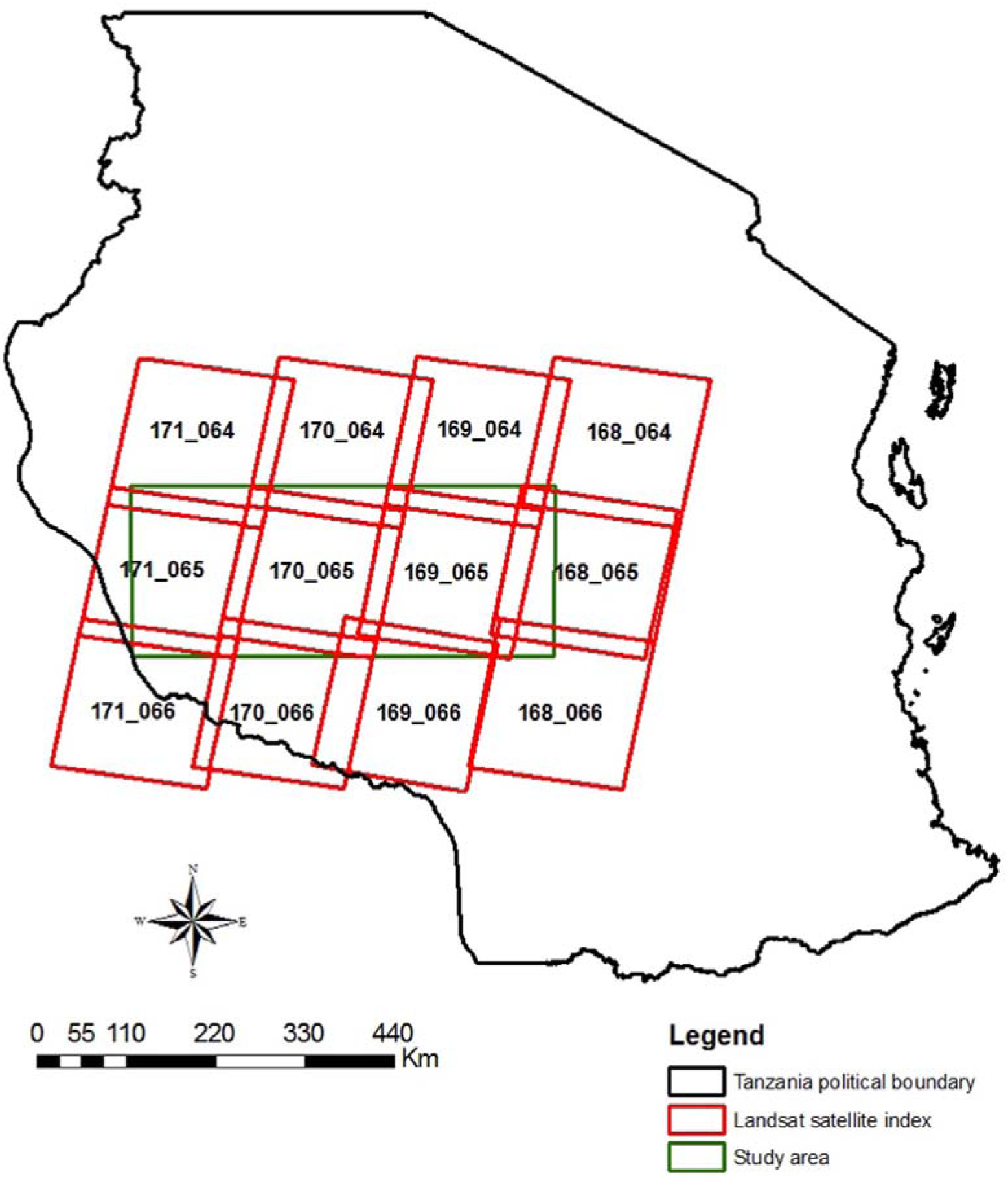

## Appendix D: Habitat loss (in Sq.km) in the Region of Interest (ROI) between the two ecosystems

| **Landuse** | **Habitat 1972** | **Habitat 1990** | **Habitat 2015** | **Change 1972-1990** | **Change 1990-2015** | **Change 1972-2015** |
| --- | --- | --- | --- | --- | --- | --- |
| Rungwa River GCA | 2494 | 2443 | 2418 | -51.5 | -24.2 | -75.8 |
| Inyonga East GCA | 3638 | 3608 | 2929 | -30.0 | -678.9 | -709.0 |
| Piti East OA | 2751 | 2732 | 1740 | -19.3 | -991.1 | -1010.4 |
| Rungwa South OA | 1828 | 1813 | 1639 | -15.2 | -173.2 | -188.4 |
| Chunya East OA | 1376 | 1367 | 1269 | -8.3 | -98.1 | -106.5 |
| Ituru Forest OA | 1328 | 1326 | 1162 | -2.4 | -163.3 | -165.8 |
| Rungwa Mzombe OA | 1945 | 1909 | 1873 | -36.6 | -35.2 | -71.8 |
| Inyonga West GCA | 1645 | 1644 | 1620 | -1.0 | -24.3 | -25.3 |
| Inyonga Central GCA | 2103 | 2106 | 1910 | 2.6 | -195.9 | -193.3 |
| Mlele North GCA | 3429 | 3343 | 3057 | -86.3 | -285.9 | -372.2 |
| Msimba East GCA | 1009 | 1007 | 1007 | -2.1 | -0.2 | -2.3 |
| Wembere South GCA | 1120 | 1108 | 822 | -11.9 | -286.0 | -297.9 |
| Rungwa North OA | 2158 | 2141 | 1106 | -17.0 | -1035.2 | -1052.1 |
| Chunya Msami OA | 1676 | 1645 | 1577 | -30.4 | -68.4 | -98.8 |
| Chunya Lukwati OA | 2318 | 2308 | 1961 | -9.4 | -347.1 | -356.4 |
|  | **30,816.53** | **30,497.68** | **26,090.49** | **(318.85)** | **(4,407.19)** | **(4,726.04)** |
| **Overall loss** |  |  |  | (318.85) | (4,407.19) | (4,726.04) |
| **Overall % loss** |  |  |  | 1.0 | 14.3 | 15.3 |

## Appendix E: Prevented habitat loss (in Sq.km) in fully protected areas in the study area

| **Landuse** | **Habitat**<br>**1972** | **Habitat**<br>**1990** | **Habitat**<br>**2015** | **Change**<br>**1972-**<br>**1990** | **Change**<br>**1990-**<br>**2015** | **Change**<br>**1972-**<br>**2015** |
| --- | --- | --- | --- | --- | --- | --- |
| Ruaha NP | 14,972.10 | 14,972.14 | 14,971.88 | 0.00 | 0.00 | 0.00 |
| Katavi NP | 4,238.81 | 4,238.81 | 4,238.81 | 0.00 | 0.00 | 0.00 |
| Lukwati GR | 3,572.98 | 3,572.96 | 3,572.97 | 0.00 | 0.00 | 0.01 |
| Rungwa GR | 8,878.05 | 8,878.05 | 8,878.01 | 0.00 | 0.00 | 0.00 |
| Muhesi GR | 1,145.15 | 1,145.13 | 1,058.74 | 0.00 | 0.00 | 86.41 |
| Lwafi GR | 2,259.84 | 2,259.83 | 2,235.13 | 0.00 | 0.00 | 24.72 |
| Kisigo GR | 5,014.72 | 5,014.71 | 5,013.43 | 0.00 | 0.00 | 0.00 |
| Rukwa GR | 4,192.60 | 4,192.56 | 4,157.98 | 0.00 | 0.00 | 34.62 |
| Piti GR | 1,687.00 | 1,769.18 | 2,072.81 | 0.00 | 0.00 | 0.00 |
|  | **45,961.25** | **46,043.37** | **46,199.76** | **-** | **0.01** | **145.77** |
|  |  |  | **%**<br>**Change** | **-** | **-** | **0.32** |

## Appendix F: Overall habitat loss (in Sk.km) in individual PA designations the entire study area

| **Landuse** | **Habitat**<br>**1972** | **Habitat**<br>**1990** | **Habitat**<br>**2015** | **Change**<br>**1972 -**<br>**1990** | **Change**<br>**1990 -**<br>**2015** | **Change**<br>**1972 -**<br>**2015** |
| --- | --- | --- | --- | --- | --- | --- |
| Rungwa River GCA | 2494 | 2443 | 2418 | -51.5 | -24.2 | -75.8 |
| Inyonga East GCA | 3638 | 3608 | 2929 | -30.0 | -678.9 | -709.0 |
| Piti East OA | 2751 | 2732 | 1740 | -19.3 | -991.1 | -1010.4 |
| Rungwa South OA | 1828 | 1813 | 1639 | -15.2 | -173.2 | -188.4 |
| Chunya East OA | 1376 | 1367 | 1269 | -8.3 | -98.1 | -106.5 |
| Ituru Forest OA | 1328 | 1326 | 1162 | -2.4 | -163.3 | -165.8 |
| Rungwa Mzombe OA | 1945 | 1909 | 1873 | -36.6 | -35.2 | -71.8 |
| Inyonga West GCA | 1645 | 1644 | 1620 | -1.0 | -24.3 | -25.3 |
| Inyonga Central GCA | 2103 | 2106 | 1910 | 2.6 | -195.9 | -193.3 |
| Lunda Nkwabi GCA | 3380 | 2638 | 2121 | -741.8 | -516.5 | -1258.3 |
| Mlele North GCA | 3429 | 3343 | 3057 | -86.3 | -285.9 | -372.2 |
| Msimba East GCA | 1009 | 1007 | 1007 | -2.1 | -0.2 | -2.3 |
| Wembere South GCA | 1120 | 1108 | 822 | -11.9 | -286.0 | -297.9 |
| Rungwa North OA | 2158 | 2141 | 1106 | -17.0 | -1035.2 | -1052.1 |
| Chunya Msami OA | 1676 | 1645 | 1577 | -30.4 | -68.4 | -98.8 |
| Chunya Lukwati OA | 2318 | 2308 | 1961 | -9.4 | -347.1 | -356.4 |
| Ruaha NP | 14,972.10 | 14,972.14 | 14,971.88 | 0.00 | 0.00 | 0.00 |
| Katavi NP | 4,238.81 | 4,238.81 | 4,238.81 | 0.00 | 0.00 | 0.00 |
| Lukwati GR | 3,572.98 | 3,572.96 | 3,572.97 | 0.00 | 0.00 | 0.01 |
| Rungwa GR | 8,878.05 | 8,878.05 | 8,878.01 | 0.00 | 0.00 | 0.00 |
| Muhsesi GR | 1,145.15 | 1,145.13 | 1,058.74 | 0.00 | 0.00 | 86.41 |
| Lwafi GR | 2,259.84 | 2,259.83 | 2,235.13 | 0.00 | 0.00 | 24.72 |
| Kisigo GR | 5,014.72 | 5,014.71 | 5,013.43 | 0.00 | 0.00 | 0.00 |
| Rukwa GR | 4,192.60 | 4,192.56 | 4,157.98 | 0.00 | 0.00 | 34.62 |
| Piti GR | 1,687.00 | 1,769.18 | 2,072.81 | 0.00 | 0.00 | 0.00 |
|  | 80,157.44 | 79,178.95 | 74,411.64 | (1,060.61) | (4,923.70) | (5,838.54) |
| Overall loss |  |  |  | (1,060.61) | (4,923.70) | (5,838.54) |
| Overall % loss |  |  |  | 1.3 | 6.1 | 7.3 |

## Appendix G: Selective logging of valuable tree species in Rungwa Game Reserve in the study area

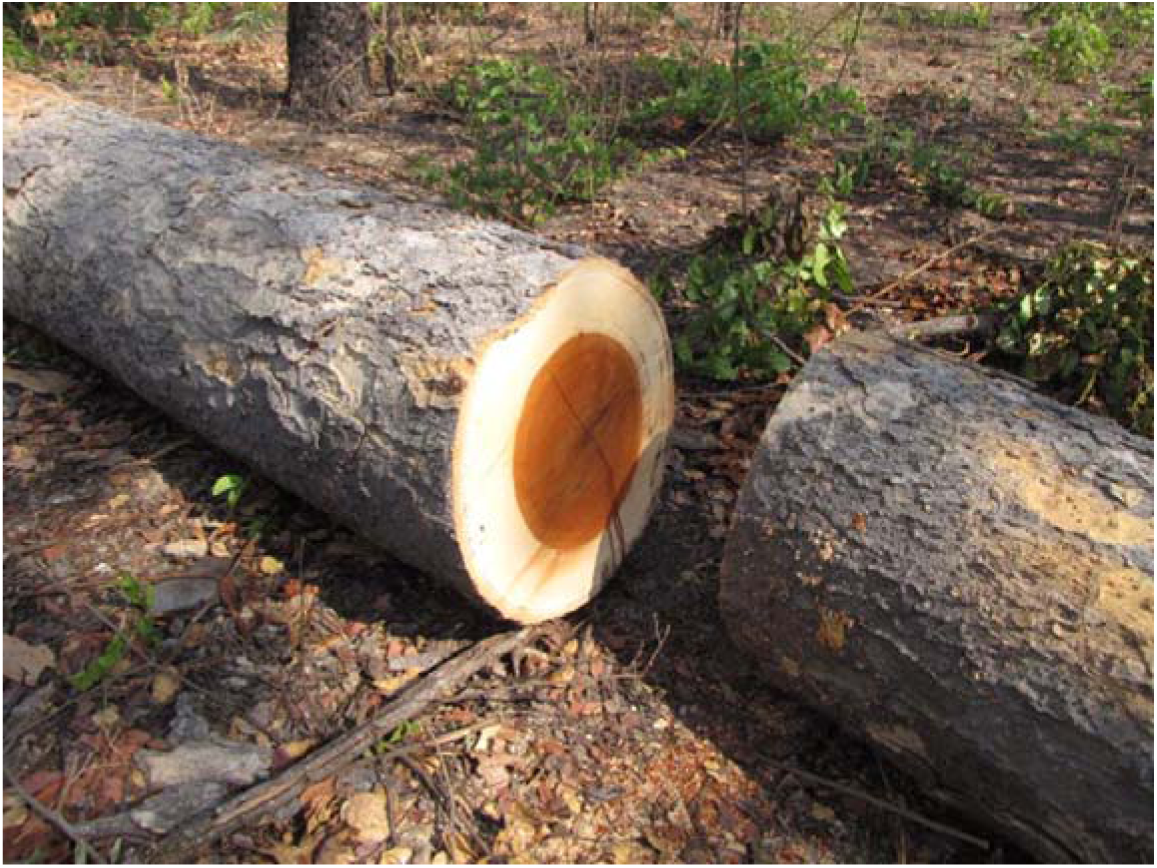

G (1) Abandoned *Afzelia quanzensis* log in Rungwa Game Reserve in the study area. Other names include pod mahogany *(spectacle case)* and East African Afzelia. Encroachment happens during wet season when most areas within the reserve are inaccessible by vehicles which is often used by game scouts during patrols. It is valued for joinery and makes attractive doors, window frames and flooring among others.

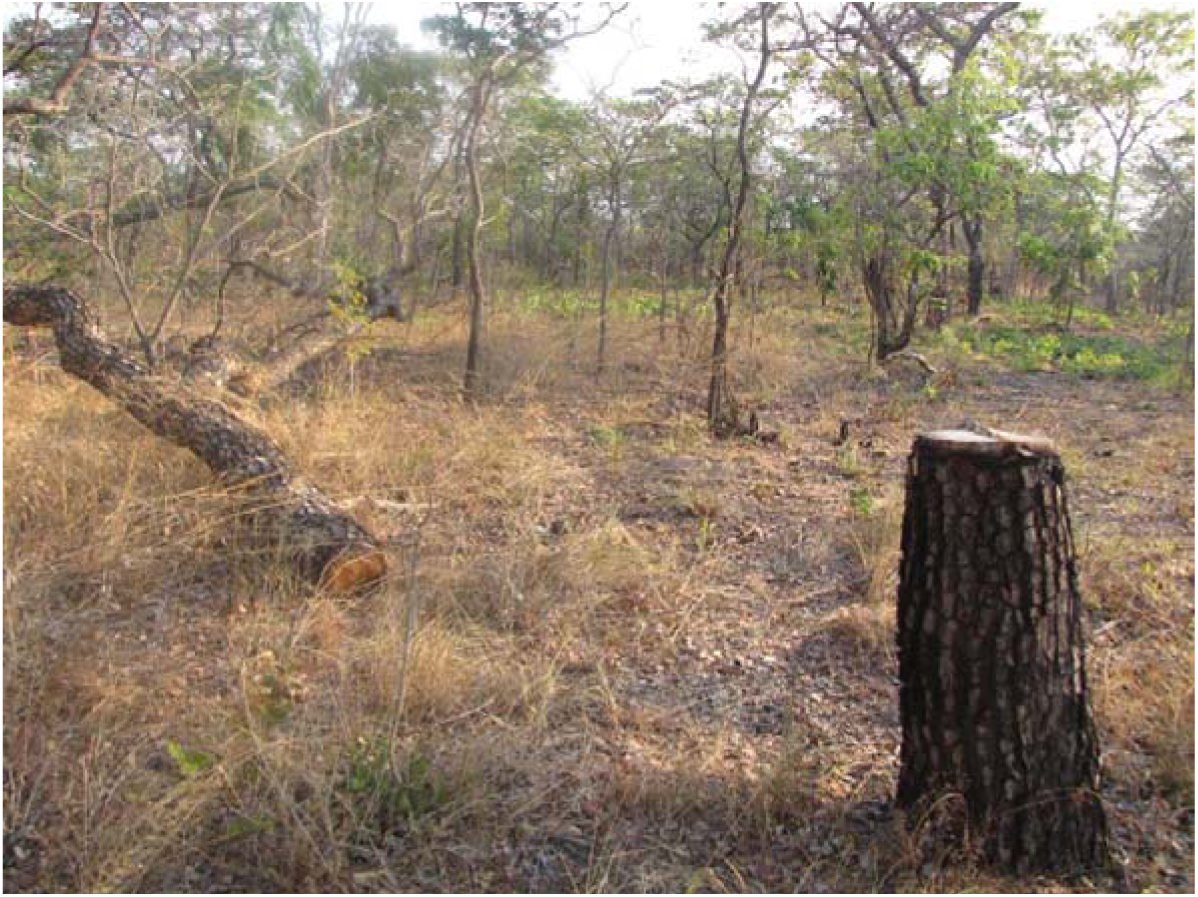

G (2) *Pterocarpus angolensis* remnant cleared for timber in Rungwa Game Reserve in the study area. It is regarded as a high valuable tree providing the highest timber quality in the market. Other names include bloodwood, paddle-wood, sealing-wax tree and wild teak.

## Appendix H: R script used to calculate change detection

~~~
**# Set working directory:**
Setwd
**#Load libraries:**
library(raster)
library(INLA)
library(spdep)
library(rgdal)
**# Reads in the raster files:**
lc_1972 <- raster("landcover1972.tif",overwrite=TRUE)
lc_1990 <- raster("landcover1990.tif",overwrite=TRUE)
lc_2015 <- raster("landcover2015.tif",overwrite=TRUE)
land_use <- raster("SA_lu_utm_36s.tif",overwrite=TRUE)
**#Crop layers to ensure they all have the same extent**
lc_1972 <- crop(lc_1972, extent(lc_1990), filename = "lc_1972_crop.tif", overwrite =TRUE)
lc_1990 <- crop(lc_1990, extent(lc_1990), filename = "lc_1990_crop.tif", overwrite =TRUE)
lc_2015 <- crop(lc_2015, extent(lc_1990), filename = "lc_2015_crop.tif", overwrite =TRUE)
**# Reprojects the rasters to the same extent and origin as roads**
lc_2015 <- projectRaster(lc_2015, roads, method = "ngb", filename -’lc2015_fmal.tif")
lc_1990 <- projectRaster(lc_1990, roads, method = "ngb", filename = "lc1990_final.tif")
lc_1972 <- projectRaster(lc_1972, roads, method = "ngb", filename = "lc1972_final.tif")
land_use <- projectRaster(land_use, roads, method = "ngb", filename = "landuse_final.tif")
**# creates the change rasters based on classes that originally were not 24 (crops) but are by the second period**
lc_change_72_90 <- writeRaster(lc_1972 != 24 & lc_1990 == 24, file = "Change_72_90.tif")
lc_change_90_15 <- writeRaster(lc_1990 != 24 & lc_2015 == 24,file = "Change_90_15.tif")
lc_change_72_15 <- writeRaster(lc_1972 != 24 & lc_2015 == 24,file = "Change_72_15.tif")
**# Calculate a change table:**
change_table_72_90 <- matrix(0, nrow = 7, ncol = 7, dimnames = list(21:27, 21:27))
change_table_90_15 <- matrix(0, nrow = 7, ncol = 7, dimnames =list(21:27, 21:27))
change_table_72_15 <- matrix(0, nrow = 7, ncol = 7, dimnames = list(21:27, 21:27)) for (i in 21:27) {for (j in 21:27) {change_table_72_90[as.character(i), as.character(j)] <- sum(values (overlay(lc_1972,lc_1990, fun=function(x,y, …) {return(x==i & y == j)})), na.rm = T) change_table_90_15[as.character(i), as.character(j)] <- sum(values (overlay(lc_1990, lc_2015, fun=function(x,y, …){return(x==i & y ==j)})), na.rm = T) change_table_72_15[as.character(i), as.character(j)] <-sum(values(overlay(lc_1972, lc_2015, fun=function(x,y, …)
{return(x==i & y == j)})), na.rm = T)}}
~~~

